# Knock-in rats expressing *Cre* and *Flp* recombinases at the *Parvalbumin* locus

**DOI:** 10.1101/386474

**Authors:** Jai Y. Yu, Jeffrey R. Pettibone, Caiying Guo, Shuqin Zhang, Thomas L. Saunders, Elizabeth D. Hughes, Wanda E. Filipiak, Michael G. Zeidler, Kevin J. Bender, Frederic Hopf, Clay N. Smyth, Viktor Kharazia, Anna Kiseleva, Thomas J. Davidson, Loren M. Frank, Joshua D. Berke

## Abstract

Rats have the ability to learn and perform sophisticated behavioral tasks, making them very useful for investigating neural circuit functions. In contrast to the extensive mouse genetic toolkit, the paucity of recombinase-expressing rat models has limited the ability to monitor and manipulate molecularly-defined neural populations in this species. Here we report the generation and validation of two knock-in rat strains expressing either *Cre* or *Flp* recombinase under the control of *Parvalbumin (Pvalb)*, a gene expressed in the critical “fast-spiking” subset of inhibitory interneurons (FSIs). These strains were generated with CRISPR-Cas9 gene editing and show highly specific and penetrant labeling of *Pvalb*-expressing neurons, as demonstrated by *in situ* hybridization and immunohistochemistry. We validated these models in both prefrontal cortex and striatum using both *ex vivo* and *in vivo* approaches, including whole-cell recording, optogenetics, extracellular physiology and photometry. Our results demonstrate the utility of these new transgenic models for a wide range of neuroscience experiments.

## Introduction

The extensive library of recombinase driver mouse models [1, 2] has enabled precise functional and structural dissection of neural circuits by providing specific and reliable genetic access to molecularly defined neural populations. When used in combination with recombinase-dependent reporter models or viruses, reporter genes can be expressed in specific neural populations [3]. For complex behavioral and large-scale *in vivo* electrophysiological experiments, rats are the model of choice. Yet their usefulness is restricted by the limited number of transgenic recombinase driver models to label molecularly defined neural populations. Thus, reliable rat recombinase driver models have great potential for accelerating progress in neuroscience.

One important application of recombinase driver models is to suppress local brain activity in real time. This can be effectively achieved by optogenetic stimulation of local inhibitory interneurons, as demonstrated in mice using inhibitory neuron-specific Cre models [2, 4]. In both cortical [5] and subcortical [6] regions, *Parvalbumin* (*Pvalb*) is expressed by GABAergic interneurons that provide perisomatic inhibition to projection neurons [7]. A rat recombinase driver model that directs transgene expression to *Pvalb*+ interneurons will therefore be a powerful tool for temporally-precise control of local circuit function.

Until recently, it has been difficult to insert exogenous genes of interest into specific genomic locations in rats, primarily due to the lack of pluripotent stem cells for *in vitro* genomic modification [8]. To overcome this limitation, genetically modified rats have been predominantly generated using Bacterial Artificial Chromosome (BAC) transgenic technology [9-11]. This involves inserting a large chunk of chromosomal DNA (200-300kb) into the genome that includes the promoter of interest, coupled with an additional coding sequence for the desired recombinase. Expression of the recombinase may then recapitulate the expression pattern of the target gene. A limited number of recombinase-driver rat strains have been created with this technique [12-14], including a transgenic with a BAC containing *Pvalb* [15].

BAC transgenics have a number of potential disadvantages, however. The BAC may or may not contain all the necessary regulatory regions to faithfully drive expression. The BAC construct is often inserted multiple times in tandem, and the insertion site cannot be controlled [16]. This can lead to overexpression or misexpression [17]. Additionally, without knowing the precise insertion locations, individual insertions responsible for the expression pattern could be gradually lost through segregation over generations of breeding.

To overcome these issues, we used the recently-developed CRISPR/Cas9-mediated gene editing [18, 19] to insert the coding sequence of recombinases into the *Pvalb* gene of Long-Evans rats. To facilitate future experiments where distinct populations of neurons are independently labelled within the same animal, we created two strains, *Pvalb^Cre^* and *Pvalb^Flpo^*, each expressing a recombinase with distinct recombination recognition sequences. Here, we validate these new models and demonstrate their utility for modulating neural networks *in vivo*.

## Results

### Molecular design

We generate two transgenic rat models expressing alternative recombinases, one expressing Cre recombinase [20] and the other expressing optimized Flp recombinase (Flpo) [21], under the control of *Pvalb* (Fig. 1A). We chose two strategies to maximize the chance of success, and both proved effective. To generate *Pvalb^Cre^*, we inserted an *Internal Ribosome Entry Site (IRES)-Cre* cassette after the termination codon of exon 5 of *Pvalb*. This modification leads to the production of a single mRNA transcript from which PVALB and CRE proteins are separately translated. To generate *Pvalb^Flpo^*, we inserted a cassette encoding the *P2A “self-cleaving” peptide* [22] followed by *Flpo* before the termination codon in exon 5. This results in the production of separate PVALB and FLPo proteins from a single ribosomal translation event.

**Figure 1.**
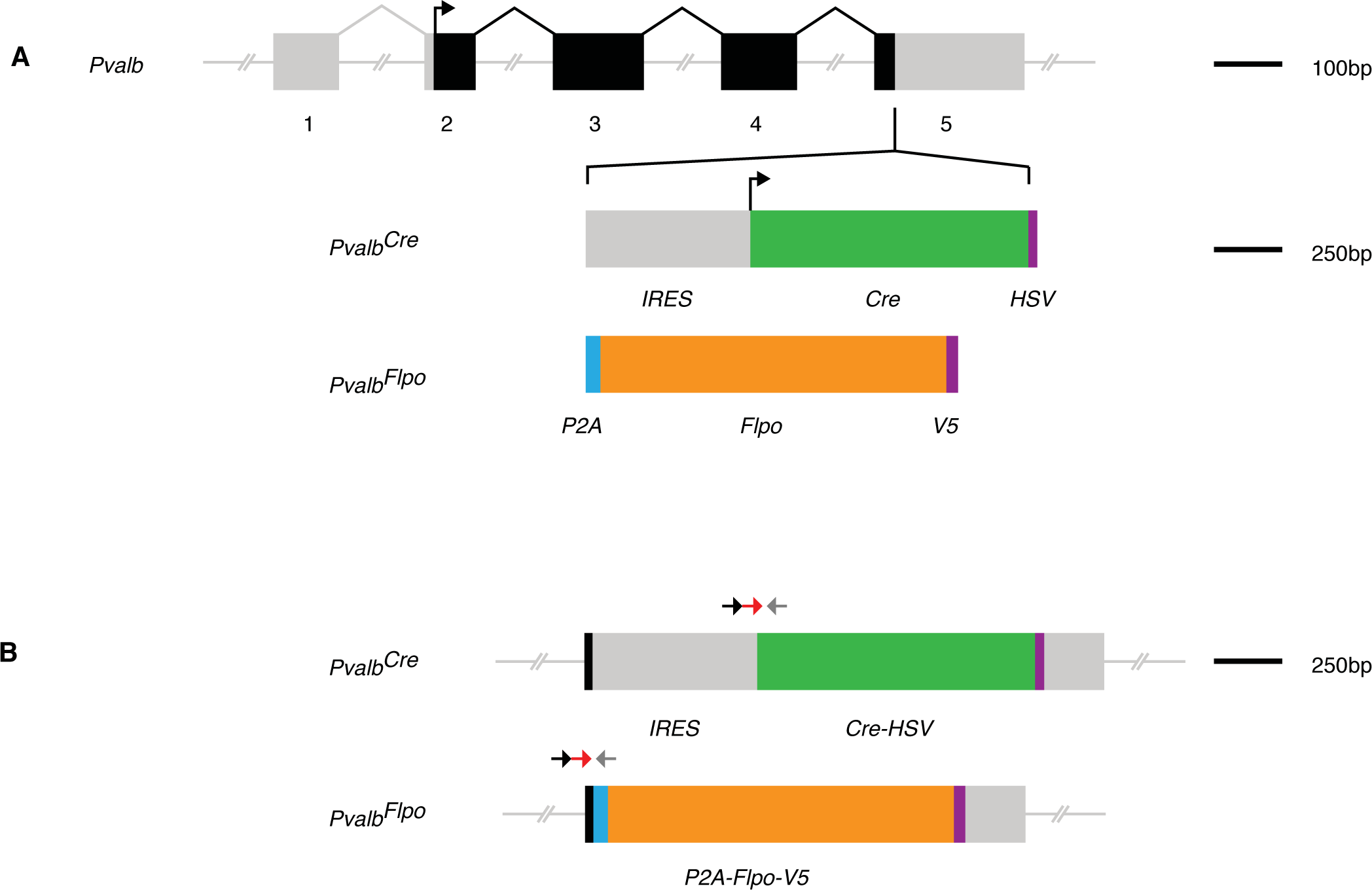
Insertion of recombinases into the *Parvalbumin* locus. **(A)** Exon structure of the *Parvalbumin* gene (gray). Arrow indicates the start of the coding region (black). Structure of the recombinase cassette inserted at the end of the coding region. IRES: internal ribosome entry site. HSV: Herpes simplex virus epitope tag. V5: V5 epitope tag. **(B)** Location of genotyping probes (arrows) in the modified locus.

To introduce the modifications, we designed single-stranded guide RNAs to induce double strand breaks near the termination codon in exon 5 of *Pvalb*. These guide RNAs were injected into zygotes together with Cas9 protein and a circular plasmid containing the gene cassette. The presence of the modifications in the offspring was detected using qPCR probes that span the joint regions (Fig. 1B).

### Consistent and specific recombinase expression in *Pvalb+* neurons

To confirm the intended recombinase expression patterns, we first used multi-color fluorescent *in situ* hybridization (FISH) to detect colocalization of recombinase and *Pvalb* mRNAs (Fig. 2) in medial prefrontal cortex (PFC) and dorsal striatum (dSTR). To further control for specificity, we included a third probe against mRNA for *Somatostatin* (*Sst*), a gene that is expressed in a largely separate population of interneurons [5].

**Figure 2.**
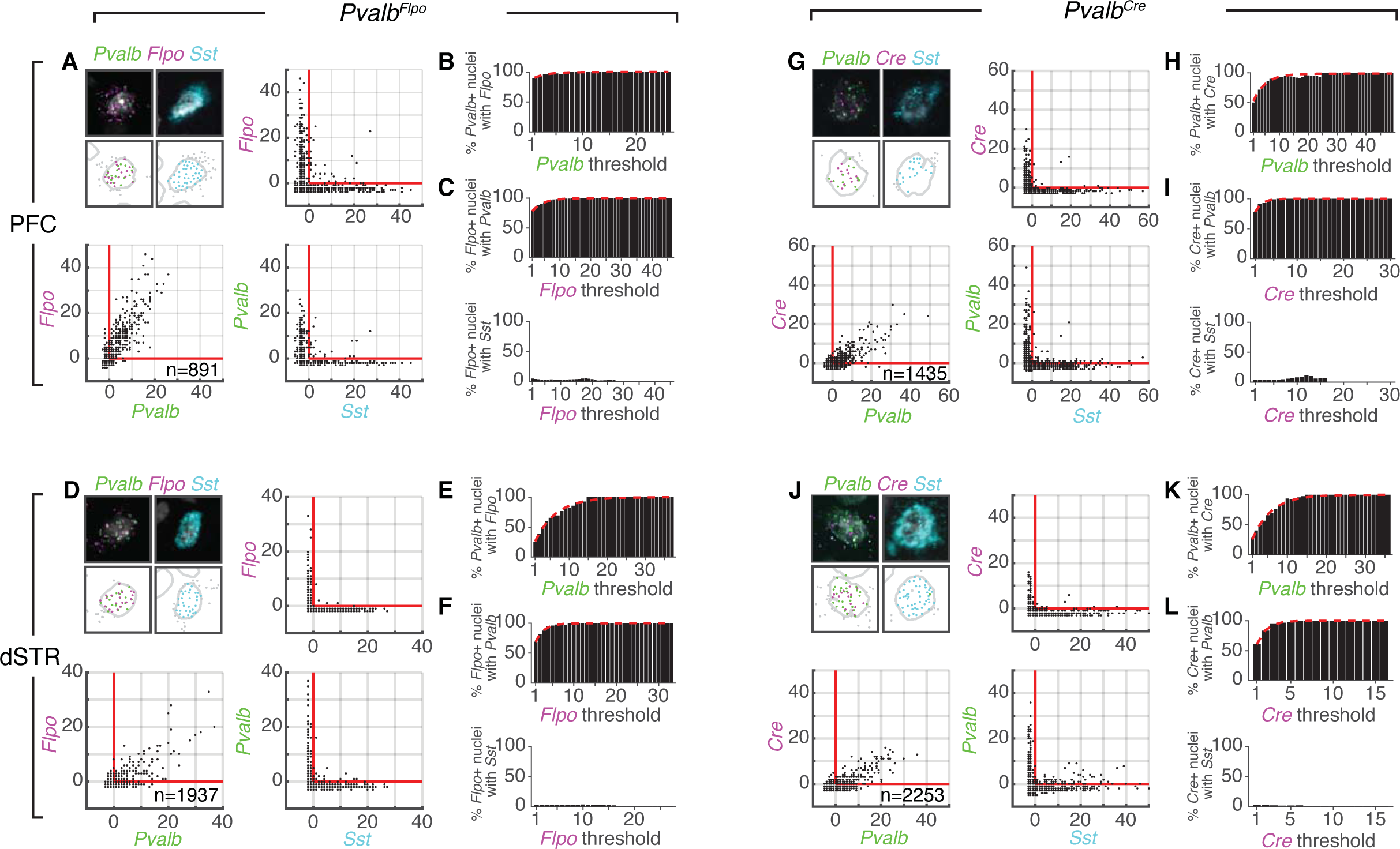
Recombinase and *Pvalb* mRNA are colocalized with high specificity and penetrance. **(A, D, G and J)** Quantification of mRNA colocalization. Example confocal images of brain sections with fluorescence *in situ* hybridization against *Pvalb* (green), *Flpo or Cre* (magenta) and *Sst* (cyan) mRNA. Nuclei are labeled with DAPI (gray). Processed images show segmentation of fluorescent mRNA puncta and nuclei. Puncta located outside of nuclei boundaries are in gray. Panel size: 50 μm. Scatter of normalized *Pvalb*, recombinase (*Flpo* or *Cre)*, and *Sst* fluorescent puncta count per nucleus. Red lines denote expected baseline count if puncta were randomly localized (see Materials and Methods). **(B, E, H and K)** Expression penetrance quantified as the percentage of nuclei with *Pvalb* puncta that had a greater than expected number of recombinase puncta. Penetrance is calculated across the range of *Pvalb* puncta thresholds (greater than or equal to abscissa). A negative exponential function (broken red line) was fitted to the distribution to estimate the asymptotic value (100% except 98.5% for (H)). **(C, F, I and L)** Expression specificity quantified as the percentage of nuclei with recombinase puncta that had a greater than expected number of *Pvalb* or *Sst* puncta. Specificity is calculated across the range of recombinase puncta thresholds (greater than or equal to abscissa) A negative exponential function (broken red line) was fitted to the distribution to estimate the asymptotic value (100%).

Manual scoring of fluorescent puncta can be subjective and difficult to reproduce reliably, especially when large numbers of tissue samples are involved. To avoid that problem, automated image processing has been successfully used to quantify FISH, especially for clinical histology samples [23, 24]. Using a similar approach, we quantified the specificity and penetrance of expression by defining nucleus boundaries, marking fluorescent mRNA puncta, and counting the number of puncta per nucleus for each fluorescent probe (Fig. 2A, D, G, and J, see Materials and Methods). From the observed number of puncta per nucleus we subtracted an estimated baseline count, to compensate for non-specific background hybridization of the fluorescent probe on the tissue sample (see Materials and Methods).

We used this normalized puncta count to quantify penetrance and specificity. We reasoned that the larger the normalized count of *Pvalb* puncta in a nucleus, the higher the confidence that it is a true *Pvalb*+ nucleus, and the larger the normalized count of recombinase puncta, the higher the confidence that a nucleus is recombinase positive. Penetrance was then defined as percentage of *Pvalb* puncta-positive nuclei with high normalized counts that also contained at least one recombinase puncta. Similarly, specificity was defined as the percentage of recombinase puncta-positive nuclei with high normalized counts that also contained at least one *Pvalb* puncta. In both cases, these values for high normalized counts were determined from the asymptote of a fit with a negative exponential function (see Materials and Methods).

For *Pvalb^Flpo^*, penetrance was close to 100% for both PFC and dSTR (Fig. 2B and E). Similarly, for *Pvalb^Cre^*, penetrance was 98.5% in PFC and close to100% in dSTR (Fig 2H and K). Specificity was similarly high, with values of close to 100% in PFC and dSTR for both *Pvalb^Flpo^* and *Pvalb^Cre^*(Fig. 2 C, F, I and L, upper panel). As expected, colocalization of either *Pvalb* or recombinase mRNA puncta with *Sst* mRNA puncta was very rare (Fig. 2 C, F, I and L, lower panel).

These very high levels of penetrance and specificity should allow effective targeting of *Pvalb*+ interneurons. Adeno-associated viruses (AAVs) encoding recombinase-dependent transgenes are widely used to achieve selective gene expression in recombinase expressing cells. We assessed the effectiveness of this strategy in our new rats. An AAV-expressing recombinase-dependent eYFP was injected into the PFC, dSTR, orbitofrontal cortex (OFC), or the dorsal hippocampus (CA1 and dentate gyrus). We quantified specificity of the AAV-mediated transgene expression approach, measured as the percentage of eYFP-expressing neurons that showed PVALB immunoreactivity. We used semiautomated image processing to mark the cell bodies in the eYFP channel. To quantify colocalization of PVALB immunoreactivity with cell bodies, we computed the mean pixel intensity value in the corresponding cell body region and compared it to the surrounding background region of the PVALB channel (Fig. 3A). We found specificity of PVALB immunoreactivity with recombinase-mediated eYFP to be 83-94% for *Pvalb^Flpo^* and 75-86% for *Pvalb^Cre^* across the sampled brain regions (Fig. 3B-C). We note that differences in specificity observed between FISH and immunohistochemistry is likely to reflect the differences in sensitivity at detecting *Pvalb* mRNA and protein, respectively, between the two methods. Overall, both FISH and immunohistochemistry demonstrate that our new models show specific and reliable recombinase expression in *Pvalb*+ neurons across multiple brain regions.

**Figure 3.**
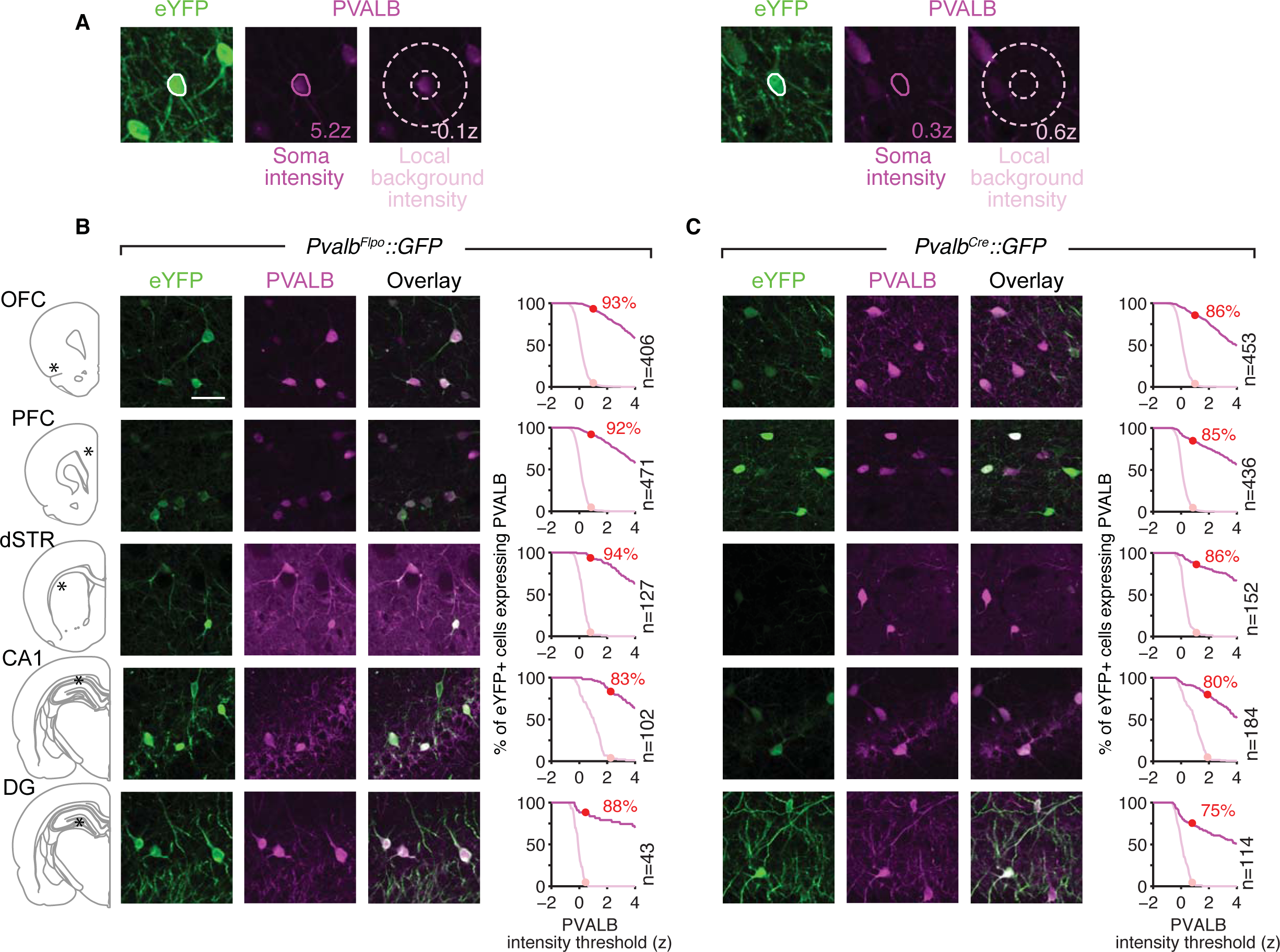
*Pvalb*-specific reporter expression mediated by recombinase-dependent AAVs. **(A)** Quantification of PVALB immunoreactivity signal. The soma of each neuron is identified from the eYFP channel. The PVALB signal intensity is the mean intensity of the corresponding area in the PVALB channel (magenta). The local background control signal (pink) is the mean intensity of a ring region around each cell (one cell width, enclosed by dotted lines). Left example shows a close up of an OFC cell with PVALB signal. Right example shows an OFC cell with a background level of PVALB signal. Intensities are normalized within each image and expressed as z scores. **(B-C)** Confocal images of immunofluorescence for eYFP (green) and PVALB (magenta) proteins across five brain regions: prefrontal cortex (PFC), orbitalfrontal cortex (OFC), dorsal striatum (dSTR), CA1 and dentate gyrus (DG). The location of each region in the brain is denoted by an asterisk in the schematic. Plots show specificity defined as the percentage of eYFP-positive soma with PVALB signal less than or equal to a given intensity. Magenta line indicates the specificity when the PVALB signal intensity is calculated from the soma. Pink line indicates the specificity expected from local background PVALB signal, which is calculated from the region surrounding each soma (see A). The light pink circle indicates upper limit of the local background PVALB signal intensity, defined as the signal intensity corresponding to 5% chance specificity. This threshold is then used to look up the true specificity estimate from the soma specificity distribution (red circle on magenta line). Number of eYFP soma counted is indicated. Recombinase-dependent AAVs expressing eYFP were injected into each region of *Pvalb^Flpo^* (B, n=4 rats) and *Pvalb^Cre^* (C, n=4 rats). Scale bar: 50 μm.

### Optogenetic activation of recombinase-expressing *Pvalb*+ neurons provides effective suppression of projection neuron activity

We functionally characterized recombinase-expressing neurons by performing whole-cell patch clamp recordings of neurons in *Pvalb^Cre^* and *Pvalb^Flpo^* rats. Using recombinase-dependent AAVs, we expressed channelrhodopsin-eYFP (ChR2-eYFP) in PFC and dSTR (Fig. 4). We first made whole-cell current-clamp recordings from fluorescent neurons. Both cortical and striatal eYFP+ cells responded to somatic current injection with high-frequency action potential firing and with sharp afterhyperpolarizations typically observed in FSIs [25, 26] (Fig. 4A-B). Full field blue light stimuli (1 ms, 470/488 nm) produced time-locked spiking activity (Fig. 4A-B), indicating that ChR2 was expressed and functional in cortical and striatal FSIs.

**Figure 4.**
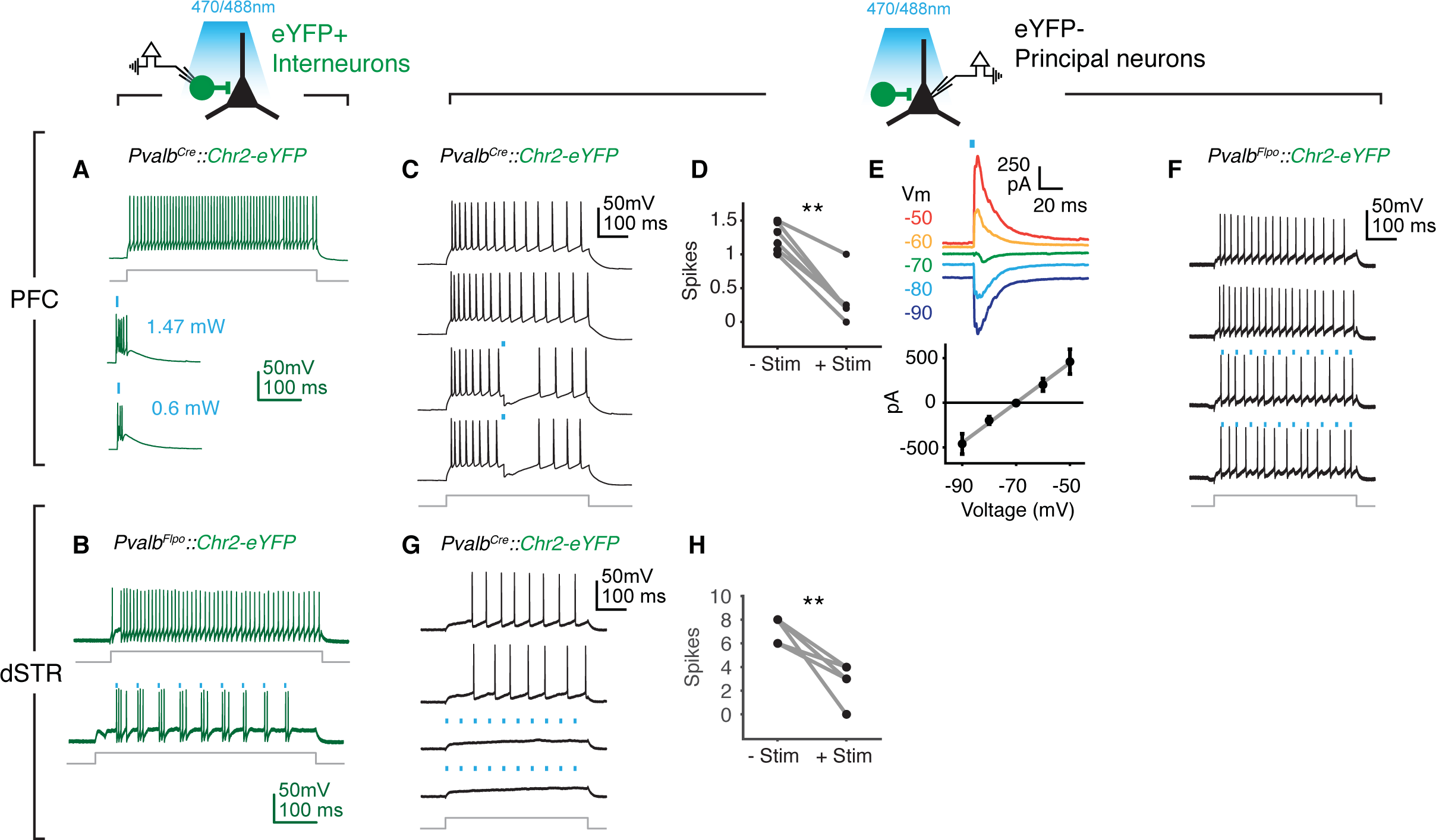
*Ex vivo* optogenetic activation of *Pvalb* expressing interneurons in PFC and dSTR. **(A)** Example response of a ChR2-eYFP-positive PFC interneuron from a *Pvalb^Cre^* rat to a depolarizing current step (400 pA, 500 ms, gray line). Example response of the PFC interneuron to different intensities of 470 nm light pulses (blue mark). **(B)** Example response of a ChR2-eYFP-positive dSTR interneuron (top trace) from a *Pvalb^Flpo^* rat to a depolarizing current step (150 pA, 500 ms, gray line). Example response of the same neuron under subthreshold current injection plus 1 ms 470 nm light pulses (bottom trace). **(C)** Example current injection (400 pA, 500 ms, gray line) traces of a PFC principal neuron from a *Pvalb^Cre^*::ChR2-eYFP rat. Top two traces show trials without light stimulation. A 1 ms light pulse (blue mark) suppressed depolarization (bottom two traces). **(D)** Quantification of mean spike count in 50 ms post stimulation or the equivalent period without stimulation for each cell (n=6). Neurons under the same stimulation condition as in (C). Wilcoxon signed rank test **p<0.05. **(E)** Inhibitory post synaptic current (IPSC) traces (top) and amplitude (bottom, mean ± SEM, n=5) during light stimulation under different holding potentials. **(F)** Example current injection (150 pA, 500 ms, gray line) of a PFC principal neuron from a *Pvalb^Flpo^*::Chr2-eYFP rat. Top two traces show trials without light stimulation. A train of 1ms light pulses (blue mark) reduced the number of evoked spikes (bottom two traces). **(G)** Example current injection (150 pA, 500 ms, gray line) traces of a dSTR neuron from a *Pvalb^Cre^* rat. A train of 1 ms light pulses (blue mark) reduced the number of evoked spikes (bottom two traces). **(H)** Quantification of spiking in the 500 ms after the onset of current injection, with or without light stimulation (n=5). Neurons under the same stimulation condition as in (G). Wilcoxon rank sum test **p<0.05.

In cortex, we saw suppression of postsynaptic excitatory neuron spiking when stimuli were delivered individually from a single or a train of light stimuli (Fig 4C, D and F). Inhibitory post synaptic current (IPSC) reversed at −70 mV, consistent with the calculated chloride reversal potential within these experimental conditions.4E). At −50 mV, peak IPSC amplitudes were 460 ± 139 pA (n = 5 cells), suggesting that multiple FSIs contributed to the resultant IPSC.

In striatum, a train of stimuli were delivered while the FSI was held to just subthreshold membrane potentials, as we subsequently found that such stimuli were necessary to suppress spiking in postsynaptic excitatory neurons (Fig. 4G and H). Consistent with this, IPSCs measured at −50 mV were smaller than those observed in cortical principal neurons (44.8 ± 5.2 pA, n = 3 cells).

We next used *Pvalb^Cre^* to optogenetically tag [27] *Pvalb*+ neurons in freely behaving rats. We expressed ChR2-eYFP in *Pvalb*+ neurons of the dSTR. An optrode array [28] was implanted to record spiking activity while blue light was delivered (Fig. 5A). In the striatum, putative *Pvalb*+ FSIs and medium spiny projection neurons (MSNs) can be distinguished by their spike waveform characteristics [29, 30]. We found neurons (n=8) that showed reliable spiking within 10 ms after light pulse onset (1 ms, 446 nm, Fig. 5B-D) and these had shorter-duration spike waveforms as expected for FSIs. Putative MSNs, identified by wide spike waveforms, did not show fast time-locked increases in spiking to light stimulation (n=39, Fig. 5G-I).

**Figure 5.**
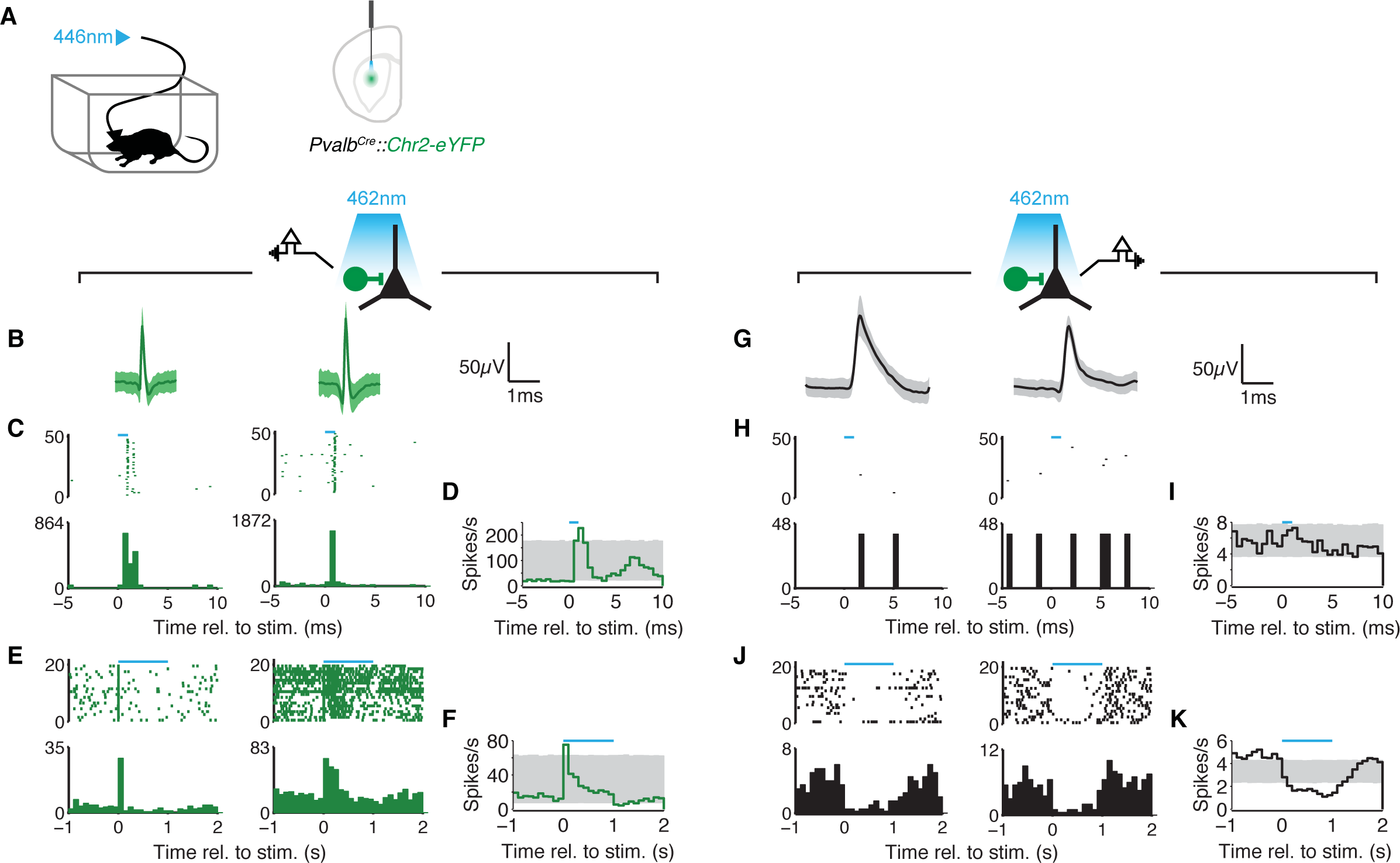
*In vivo* optogenetic activation of *Pvalb*+ interneurons suppresses principal neuron activity in dSTR. (A) AAV-DIO-ChR2-eYFP was injected into the dSTR of *Pvalb^Cre^* rats (n=2). An optrode array was used to deliver 446 nm light while recording single unit activity. **(B-F)** Spiking activity of ChR2+ dSTR interneurons. **(B)** Spike waveforms (mean ± SD) for two example tagged ChR2+ neurons. **(C)** Raster and histogram of activity of the same example tagged neurons aligned to 1 ms stimulation (blue bar). **(D)** Mean histogram of activity of multiple interneurons (n=8) aligned to 1 ms stimulation. Gray indicates 95% CI of time bin permutation control (n=10000). **(E)** Raster and histogram of activity of the example interneurons aligned to 1 s stimulation (blue bar). **(F)** Mean histogram of activity of multiple interneurons (n=5) aligned to 1 s stimulation. Gray indicates 95% CI of time bin permutation control (n=10000). **(G-K)** Spiking activity of dSTR principal neurons. **(G)** Spike waveform (mean ± SD) of 2 example principal neurons. **(H)** Raster and histogram of activity of the example neurons aligned to 1 ms stimulation (blue bar). **(I)** Mean histogram of activity of multiple principal neurons (n=39) aligned to 1 ms stimulation. Gray indicates 95% CI of time bin permutation control (n=10000). **(J)** Raster and histogram of activity of the example neurons aligned to 1 s stimulation (blue bar). **(K)** Mean histogram of activity of multiple principal neurons (n=36) aligned to 1 s stimulation. Gray indicates 95% CI of time bin permutation control (n=10000).

Longer (1s) stimulation produced both transient excitation in putative FSIs (n=5, Fig. 5E-F) and marked suppression of putative MSN firing (n=36, Fig. 5J-K). Such FSI-induced MSN inhibition is consistent with prior [31] and current *ex vivo* results, but stands in contrast to prior *in vivo* studies, which found little evidence that spontaneous FSI spikes affect MSN spiking [30, 32]. One plausible explanation is that FSI inhibition of MSNs *in vivo* requires synchronized spiking of multiple FSIs, as produced by optogenetic activation under our experimental conditions.

### Behavioral state-dependent activity of *Pvalb*+ neural populations

Finally, we asked whether fiber photometry could be used to examine population activity of *Pvalb*+ neurons in unrestrained animals. Using a Flp-dependent AAV in *Pvalb^Flpo^* rats, we expressed the calcium indicator GCaMP6f in *Pvalb*+ neurons of PFC and recorded calcium-dependent fluorescence through implanted optical fibers (Fig. 6A, 9 recording sessions across 3 animals). Each animal was monitored while freely behaving in its home cage for 3 hours.

**Figure 6.**
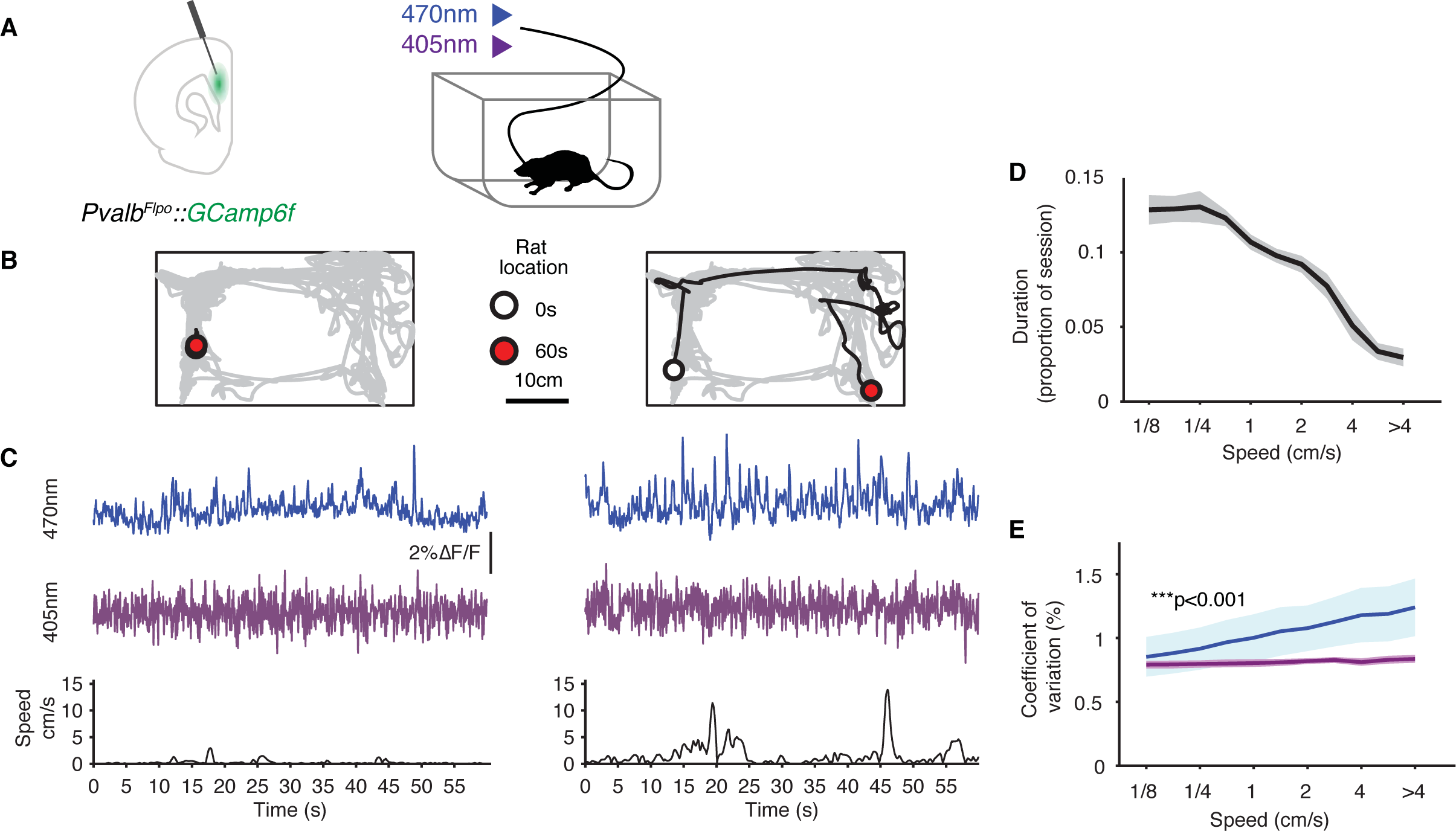
Behavior-dependent calcium signals in *Pvalb* expressing neural populations. **(A)** GCaMP6f expressed in *Pvalb*+ neurons of PFC of *Pvalb^Flpo^* rats (n=3). GCaMP6f emission from 470 nm (calcium-dependent) and 405 nm (isosbestic control) excitation were collected. **(B)** Example rat movement trajectory in the home cage over 60 s intervals (black) overlaid on the movement trajectory for the entire session (gray). The white and red circles indicate the start and end locations respectively. **(C)** GCaMP6f emission from 470 nm (blue) and emission from 405 nm excitation (purple). Signal traces are shown as percentage deviation from the background normalized mean. Speed for the corresponding period is indicated below the signal traces. These examples illustrate contrasting calcium dependent GCaMP6f signal fluctuations between periods with distinct movement patterns. **(D)** Proportion of 5 s intervals spent at different speeds for each recording session (mean ± SEM). 9 sessions were included (rat 1, 3 days; rat 2, 2 days and rat 3, 4 days). **(E)** Fluctuation in photometry signal amplitude as a function of movement speed in 5 s intervals (mean ± SEM) for GCaMP6f emission from 470 nm (blue) and 405 nm (purple) excitation. Signal fluctuation is expressed as the coefficient of variation, which is the standard deviation divided by mean of the signal. Wilcoxon rank sum test for a difference in slopes: ***p<0.001.

We compared *Pvalb*+ population activity during periods with distinct movement patterns (Fig. 6B). During periods of low movement speeds, fluctuations in calcium-dependent GCaMP6f fluorescence tended to be small (Fig. 6B-C, left). In contrast, during periods of higher movement speeds, we saw larger fluctuations in calcium dependent GCaMP6f fluorescence (Fig. 6B-C, left). To describe the relationship between movement speed and GCaMP6f fluorescence, we first characterized the proportion of time the rat spent at a range of movement speeds in a given session (Fig. 6D). We next quantified the fluctuation in GCaMP6f activity during periods at these movement speeds by computing the coefficient of variation (CV), which is the standard deviation of the signal divided by its mean. We found GCaMP6f signal CV was smallest during low speed periods, and increased with higher movement speeds (Fig. 6E). The relationship between CV and speed was significantly stronger for the 470 nm calcium-dependent fluorescence compared with the 405 nm isosbestic control fluorescence. These findings suggest an increase in *Pvalb*+ population activity with speed, complementing previous findings of speed-dependent *Pvalb*+ firing in visual cortex [33]. These results demonstrate that *in vivo* fiber photometry can be used to monitor population of *Pvalb*+ neurons using our recombinase model to examine behavioral state-dependent activity patterns.

## Discussion

We created two knock-in recombinase rat models that enable the labeling of *Pvalb*+ interneurons with high penetrance and specificity. The two alternative *Pvalb* recombinase driver models enable the labeling of multiple specific neural populations within the same animal. As one way to achieve this, *Pvalb^Flpo^* animals can be bred with Cre-expressing rat strains [12-14, 34, 35]. As another example, retrogradely-traveling Cre-dependent viruses [36, 37] can be used in *Pvalb^Flpo^* animals to label specific projection pathways between two brain regions, while Flp-dependent constructs can be used at the same time to label *Pvalb*+ neurons. Finally, the *Pvalb^Flpo^* animals are suitable for intersectional genetic methods [2, 38], using viral constructs that require both Flpo and Cre to permit reporter expression. By crossing *Pvalb^Flpo^* with a strain that expresses *Cre* in a neural population that has overlap with *Pvalb*, the specific subset that is double positive for both *Cre* and *Flpo* can be selectively labeled.

One potential side effect of CRISPR/Cas9-mediate gene editing is the introduction of off-target mutations in the genome [39]. We are minimizing the potential effects of these off-target changes by backcrossing each generation to wildtype animals and maintaining the modified strains as heterozygotes. All of our experiments were performed on animals backcrossed for a minimum of 3 generations. Any potential off-target changes should be progressively eliminated over successive generations of backcrossing.

These new knock-in *Pvalb* recombinase models expand the current genetic toolkit available in rats. The genetic access afforded by these models to a molecularly-defined interneuron population will complement the ability to perform large scale electrophysiology recordings and complex behavioral tasks that are feasible in rats. Together, they enable new classes of experiments to investigate the structure and function of neural circuits in controlling behavior.

## Author contributions

CG, SZ and JYY designed the *Pvalb^Cre^* construct. CG and SZ generated the *Pvalb^Cre^* model. TLS, JDB and JYY designed the *Pvalb^Flpo^* construct. TLS, EDH, WEF and MGZ generated the *Pvalb^Flpo^* model. KJB and FH performed the *ex vivo* electrophysiology experiments. JRP performed the FISH and *in vivo* electrophysiology experiments. CNS and JYY performed viral injections for histological characterization, and the photometry experiments with TJD. VK performed the histology. AK maintained the transgenic colonies. JYY analyzed the FISH, immunofluorescence, and photometry data, and electrophysiology data with JRP. JYY, JDB and LMF conceived and designed the project, and wrote the manuscript with input from all authors. JDB and LMF directed the project.

## Acknowledgements

We thank Lief Fenno, Joanna Mattis and Charu Ramakrishnan for generously sharing reagents for proof of concept experiments, and Duda Kvitsiani, Hongkui Zeng and Alla Karpova for advice on the feasibility of the project. We also thank Alla Karpova and Carlos Brody for their financial support for generating the *Pvalb*^*Cre*^ model.

## Materials and Methods

Animals were housed in AAALAC-accredited facilities in accordance with the National Research Council’s guide for the care and use of laboratory animals. Procedures were approved by the University of Michigan’s, the University of California San Francisco’s and Janelia Farm Research Campus’ Institutional Animal Care & Use Committees.

### Knock-in rat generation

CRISPR/Cas9 technology [18, 19] was used to generate genetically modified rat strains to express the *Flpo* [21] or *Cre* transgene under the control of *Pvalb*.

### Locus targeting

#### Pvalb^Cre^

The donor construct contains two homology arms that were PCR amplified from the *Pvalb* gene using Long-Evans rat genomic DNA. The 1167 bp 5’ arm was amplified with primers 5’ TCT ACT AGT GGG AAA GGG ATA AAC CAG G 3’ and 5’ GTC TTA ATT AAT TAG CTT TCG GCC ACC AGA G 3’. The 1693 bp 3’ arm was amplified with primers 5’ CAT AGC GCT GCG CTG ACT GCT TGG GTC TC 3’ and 5’ CAT AGC GCT GCG CTG ACT GCT TGG GTC TC 3’. The PCR products were cloned into a pBluscript vector containing an *Encephalomyocarditis Internal Ribosome Entry Site (IRES)-Cre* cassette.

The single guide RNA (sgRNA) 5’ TCT GGT GGC CGA AAG CTA AG 3’ was designed to target and generate a double strand break at the 3’ terminus of the coding region of exon 5. It was obtained by *in vitro* transcription using a template of PCR product primers 5’ CCT TAA TAC GAC TCA CTA TAG GTC TGG TGG CCG AAA GCT AAG GTT TTA GAG CTA GAA ATA GC 3’ and 5’ AAA AGC ACC GAC TCG GTG CCA CTT TTT CAA GTT GAT AAC GGA CTA GCC TTA TTT TAA CTT GCT ATT TCT AGC TCT AAA AC 3’. The gRNA was tested *in vitro* by digesting a 520 bp PCR products with the gRNA and CAS9 protein.

### Zygote microinjection

7-week-old female Long-Evans rats (Charles River Cat .006) were used as embryo donors. They were superovulated with equine chorionic gonadotropin (30 units) and human chorionic gonadotropin (30 units) 48 hours apart and mated with studs. 1-cell zygotes were collected the next morning. A mix of CAS9/gRNA/circular targeting vector (100 ng/100 ng/20 ng) was microinjected into the zygotes. The injected zygotes were transferred into pseudo pregnant Sprague Dawley rats. Fourteen F0 pups were generated. Two F0 rats contained the correct knock-in cassette and were bred with wildtype Long-Evans rats.

### Screening

The F0 pups were screened by nested PCR using primers outside the homology arms (F1, F2, R3 and R4) with primers inside the *IRES-Cre* cassette (R1, R2, F3 and F4). F1: 5’ TGG GAA TCA CCT GTG TAC CA 3’. F2: 5’ CAC CTG AAG AGT GTT ATG CC 3’. F3: 5’ TAC CGG AGA TCA TGC AAG CT 3’. F4: 5’ ATC CGT AAC CTG GAT AGT GAA 3’. R1: 5’ AGG AAC TGC TTC CTT CAC GA 3’. R2: 5’ CCT AGG AAT GCT CGT CAA GA 3’. R3: 5’ ACC TCA TGG TCT AAG TGG GA 3’. R4: 5’ AGT GGT GCA CAC CCT GAT AC 3’

#### Pvalb^Flpo^

A double-stranded DNA plasmid donor was synthesized (Invitrogen GeneArt, Thermo Fisher Scientific) to introduce the following elements before the termination codon in exon 5: a glycine-serine-serine linker with porcine teschovirus-1 self-cleaving peptide 2A (P2A) [22], followed by *Flpo* recombinase [21] and the V5 peptide tag (GKPIPNPLLGLDST) [40]. To mediate homologous recombination with the chromosome, a 5’ arm of homology (794 bp of genomic DNA 5’ to codon 110) and a 3’ arm of homology (469 bp of genomic DNA downstream of the termination codon) were used.

Two guide RNA (sgRNA) targets and protospacer adjacent motifs (PAM) were identified downstream of rat *Pvalb* termination codon (ENSRNOG00000006471) using published algorithms [41]. The sgRNA targets were cloned into plasmid pX330 (Addgene.org plasmid #42230, a kind gift of Feng Zhang) as described [42]. The guide targets were C9G1: 5’ TCT GGT GGC CGA AAG CTA AG 3’ PAM TGG and C9G2: 5’ AGT CAG CGC CAC TTA GCT TT 3’ PAM: CGG. Circular pX330 plasmids were co-electroporated into rat embryonic fibroblasts with a PGKpuro plasmid [43]. Genomic DNA was prepared from cells that survived transient selection with puromycin (2μg/ml). A 1125 bp DNA fragment spanning the expected Cas9 cut site was PCR amplified with forward primer 5’ CTG GAT CCC TCC CAC ACA GA 3’ and reverse primer 5’ TGG TCC TTC GCT CTC TCT CA 3’. Amplicons were subjected to CEL I endonuclease digestion essentially as described [44]. Briefly, DNA amplicons were melted and re-annealed, then subjected to CEL I digestion. The digested DNA fragments were separated by agarose gel electrophoresis. Gels were stained with Sybr Gold (Invitrogen S11494) to detect insertion/deletion mutations (indels) in the amplicons. The presence of indels produced by non-homologous endjoining repair of induced double strand breaks resulted in the presence of lower molecular weight DNA fragments following CEL I digestion for C9G1 but not C9G2 sgRNA. sgRNA C9G1 was chosen for rat zygote microinjection. The C9G1 sequence was interrupted by the insertion of the synthetic sequence, thus preventing Cas9 cleavage of the chromosome after correct insertion of the DNA donor.

### Zygote Microinjection

Rat zygote microinjection was carried out at described [45]. Circular pX330 plasmid DNA containing C9G1 sgRNA was purified with an endotoxin free kit (Qiagen, Germantown, Maryland). Circular plasmid donors were similarly purified. Zygotes were microinjected with a solution containing a C9G1 pX330 plasmid (5 ng/μl) [46] and 10 ng/μl of plasmid of the circular DNA donor plasmid in 10 mM Tris, pH 7.4, 0.1 mM EDTA. To ensure that the plasmid DNA mixtures did not cause zygote death or block development to the blastocyst stage, fertilized mouse eggs were microinjected with the nucleic acid mixture and cultured to the blastocyst state.

Rat zygotes for microinjection were obtained by mating Long Evans female rats, superovulated with equine chorionic gonadotropin (20 units) and human chorionic gonadotropin (50 units), with Long Evans male rats from an in-house breeding colony. A total of 693 rat zygotes were microinjected, of which 523 survived injection and were transferred to pseudopregnant female rats.

Genomic DNA from the 118 rat pups born was purified from tail tip biopsies with a Qiagen DNeasy kit. In the first pass genotyping screen used PCR primers specific for the *Flpo* coding sequence (forward primer 5’ TGA GCT TCG ACA TCG TGA AC 3’, reverse primer 5’ TGC TGG TGT ACT CGA AGC TG 3’). Thirteen of the pups were positive for integration of the *Flpo* transgene. These potential founders were then screened for correct 5’ integration with a forward primer lying outside of the 5’ arm of homology (5’ ATT ACA GCC CTG TCC TTT ACT TTT TAT 3’) and a primer lying inside the *Flpo* sequence (5’ GAT GAT GGT GTT GTA GCT CAT GAA GG 3’). Similarly, a primer lying outside of the 3’ arm of homology (5’ GCT CAC TCT TGT GGG TAT TTA TGA C 3’) and a primer internal to *Flpo* (5’ ACT ACT TCG CCC TGG TGT CCA GGT ACT A 3’) were used to identify founders with correct integration on the 3’ side *Pvalb*. PCR amplicons were sequenced to verify correct integration. Primers internal to *Flpo* were used to verify the *Flpo* coding sequence. Correct integration was found in eight of the thirteen potential founders. Germline transmission of the *Flpo* knock-in rats was confirmed in the offspring of one founder and a line of *Pvalb^Flpo^* knock-in rats was established. The *Pvalb^Flpo^* knock-in G0 founder rats were mated with Long Evans rats. The G1 offspring were screened with the junction spanning primers and sequenced to confirm transmission of the correct knock-in sequence.

### Colony management

The line was maintained by outcrossing males from each generation with wildtype females (Charles River) for a minimum of 3 generations. The insertion was verified by sequencing. Offspring were genotyped using real-time PCR (Transnetyx). *Pvalb^Cre^* primers are located at the *Ires-Cre* junction (Fwd. Primer: 5’ TTC CTT TGA AAA ACA CGA TGA TAA TAT GGC 3’. Rev. Primer: 5’ GGT AAT GCA GGC AAA TTT TGG TGT A 3’. Reporter 1: 5’ ACA ACC ATG TCC AAT TTA 3’).

*Pvalb^Flpo^* primers are located at the junction of the *Pvalb-P2A-Flpo* cassette (Fwd. Primer: 5’ CAA GCT GGT GCA GAA CAA GTA C 3’, Rev. Primer: 5’ TGG ACA CGC TTG TCT TGG T 3’ and Reporter 1: 5’ CTG GGC GTG ATC ATT C 3’).

### Virus injection

Virus (250-500 nl) was injected at a rate of 0.2 μl/minute using a syringe with a beveled 33G needle (Hamilton Nanofil 10 μl). The syringe was controlled with a stereotax mounted syringe pump (kd Scientific). After each injection, the needle was left in place for 3-10 minutes before retraction to ensure diffusion of the virus. The virus was allowed to express for 4-5 weeks.

The following viruses were used: AAV DJ ef1α fDIO-eYFP-WPRE, titer 2.4 x 10^13^, Stanford Virus Core; AAV2/5 ef1α fDIO-ChR2(H134R)-eYFP, titer 1.2×10^12^, UNC Virus Core; AAV DJ Syn FRT-rev GCamp6f, titer 1.58×10^13^, Janelia Farm Virus Core; AAV DJ ef1 DIO-eYFP, titer 8.9 x 10^13^, Stanford Virus Core and AAV2/5 CAG DIO-ChR2(H134R)-eYFP, titer 10^13^, Virovek.

### Stereotactic coordinates

PFC: AP +2.5, ML ±1.5 and DV −2.5 at 20º relative to vertical midline. OFC: AP +3.6, ML ±3.4 and DV −4. Dorsal hippocampus: AP −3.6, ML ±2.6 and DV −2.7. Dorsal striatum: AP +0.8, ML ±3.8 and DV −4.3.

### Immunofluorescence

Animals were deeply anesthetized with Euthasol and intracardially perfused with heparinized phosphate-buffered saline (PBS), followed by 4% paraformaldehyde in PBS, pH 7.4. Brains were removed, post-fixed in the same fixative for 2 hours at 4 ºC and transferred to PBS. On the following day, brains were transferred into 30% sucrose and stored for 5 days at 4 ºC. Frozen 50μm-thick coronal sections were cut on a cryostat (Microm, Thermo Fisher Scientific), collected in 24-well plates and stored in PBS at 4 ºC. Free-floating sections containing regions of interest: PFC, OFC, dStr or the dorsal hippocampus (CA1 and dentate gyrus) were transferred into a 12-well plate in PBS, permeabilized with 50% ethanol for 10 minutes, and rinsed in PBS. Sections were then blocked with 10% normal donkey serum in PBS for 30 minutes and incubated for 48 hours at 4 °C on an orbital shaker with a mixture of primary antibodies: rabbit polyclonal anti-GFP (ab290, Abcam, 1:10000) and mouse monoclonal anti-PVALB (Sigma-Aldrich, 1:1000) diluted in PBS containing 0.05% Triton X-100. Next, sections were washed in PBS, incubated with 2% normal donkey serum for 10 min, and incubated for 4 hours with the secondary antibodies: Alexa Fluor 488-labeled donkey anti-rabbit and Alexa Fluor 594 or 647-labeled donkey anti-mouse antibodies (Thermo Fisher Scientific, 1:300). After staining, sections were rinsed in PBS and coverslipped using ProLong Gold Antifade mountant medium containing DAPI (Thermo Fisher Scientific).

### mRNA fluorescence *in situ* hybridization

Brains were removed from animals following deep anesthesia under 5% isofluorane inhalation, submerged in Optimal Cutting Temperature cutting medium (Tissue-Tek) and placed on powdered dry ice. Frozen brains were stored in an airtight container at −80 ºC (overnight to 2 weeks). The brains were sectioned at 20 μm on a cryostat and mounted on glass slides (SuperFrost Plus, Fisher Scientific). Sections were fixed in 4% PFA at 4 ºC for 15 minutes and dehydrated through 50%, 75%, 100% and fresh 100% ethanol at RT for 5 minutes each. Slides were dried completely for 5 minutes. A hydrophobic barrier (Advanced Cell Diagnostics) was drawn around each section. Slides were rinsed twice in 1xPBS (~1-3 minutes) and incubated with Protease IV reagent (30 minutes at RT). Fluorescent probes (RNAScope©, Advanced Cell Diagnostics, *Pvalb* Cat. 407821-C2 and *Sst* Cat. 412181-C3 with Cre Cat. 312281 or *Flpo* Cat. 448191) were added each slide (2 hours, 40 ºC) followed by manufacturer specified washing and amplification. DAPI was added to the slides before mounting (Prolong Gold, Thermo Fisher Scientific).

### Imaging

Immunofluorescence and fluorescent *in situ* hybridization images were taken with a Nikon spinning disc confocal microscope using manufacture recommended filter configurations with a 20X objective (Plan Apo Lambda NA 0.75 and image area 665.8×665.8 μm) for fluorescence immunohistochemistry or a 40X objective (Plan Apo Lambda NA 0.95 and image area 332.8×332.8 μm) for fluorescent *in situ* hybridization. Images were acquired at 2048×2048 pixels at 16-bit depth.

### Image quantification

#### mRNA fluorescent *in situ* hybridization

We used the MIPAR image analysis software (www.mipar.us) to segment cell boundaries and fluorescent puncta using separate processing pipelines.

The DAPI channel of each image was used to define nucleus boundaries, which were segmented as follows. The image was first histogram equalized to compensate for uneven illumination (512×512 pixel tiles) and convolved with a pixel-wise adaptive low-pass Wiener filter (5×5 pixel neighborhood size) to reduce noise. The image was then contrast adjusted to saturate the top and bottom 1% of intensities. Bright pixels were segmented as objects using an adaptive threshold, defined as the pixel intensity greater than 110% of mean in the surrounding 30-pixel window. Image erosion followed by dilation was used to reduce noise (5-pixel connectivity threshold, 10 iterations). The Watershed algorithm was applied to improve object separation. Objects larger than 6000 pixels (i.e. clustered nuclei) were identified and reprocessed to improve separation within the cluster. Since mRNA fluorescent puncta can be located in the endoplasmic reticulum surrounding the nuclei, we dilated the boundaries of each segmented nuclei by 5 pixels to include these regions.

For segmenting fluorescent puncta, we used the following steps. To reduce noise and improve contrast, images were filtered using a Top-hat filter (15-pixel radius), Wiener filter (99×99 pixel neighborhood size) followed by contrast adjustment to saturate the top and bottom 1% of intensities. Bright regions were segmented using the extended-maxima transform (8-connected neighborhood, 5 H-maxima). A Watershed algorithm followed by erosion was used to improve object separation. The location of each puncta is defined as the centroid of the segmented object. This was done individually for each of the three fluorescent probe image channels (*Pvalb*, recombinase and *Sst*).

For each fluorescent probe image channel, we counted the number of segmented puncta objects lying within a nucleus’ boundary. Since there can be puncta located outside of nuclei from non-specific fluorescence probe hybridization, we needed to estimate the number of puncta expected per nucleus by chance due to this background. We computed the expected number of puncta within a nucleus’ boundary by randomly relocating (spatially permuting) all identified puncta in an image channel, either lying within or outside of nuclei. We repeated the permutation 1000 times for each image. For each permuted image, we again counted the number of puncta lying within a nucleus’ boundary. We created a distribution of expected puncta per nucleus count from the permuted images. We then used a p < 0.005 threshold (99.5th percentile) to estimate the expected puncta count per nucleus by chance. We subtracted the baseline puncta count from the observed puncta count to derive the normalized puncta count per nucleus. We note this is a conservative approach for estimating the baseline levels since the null assumption is that all puncta are due to spurious background hybridization.

To estimate penetrance, we first calculated the percentage of *Pvalb+* nuclei that also had greater-than-background counts of *recombinase* puncta. We used a range of *Pvalb* puncta counts to define *Pvalb+* nuclei, rather than a single, arbitrary threshold that could be readily affected by experimental conditions. As the puncta count increased, a nucleus became increasingly certain to be *Pvalb+*. We reasoned that the true penetrance likely can be estimated from the asymptotic value across the range. We fitted a negative exponential function α(1-*e*^β(-x+γ)^) and defined the penetrance as α, which corresponds to the asymptotic value on the y axis.

To estimate specificity, we first calculated the percentage of nuclei with recombinase puncta counts that also had greater than chance counts of *Pvalb* puncta, for a range of *recombinase* puncta counts. We then used the same method as the penetrance estimation to determine the asymptotic value on the y axis for the negative exponential function fit.

### Immunofluorescence

Neuron cell bodies were identified from the eYFP channel as high signal intensity regions. This was done using the following functions from the MATLAB Image Processing Toolbox. To compensate for uneven background illumination, each image was first histogram normalized (20×20 pixel tiles). A pixel-wise adaptive low-pass Wiener filter (5×5 pixel neighborhood size) followed by a Median filter (3×3 pixel neighborhood size) was used to reduce noise. The preprocessed image was thresholded using Otsu’s method to isolate bright pixels regions. Regions were smoothed with morphological opening using a 5-pixel disk-shaped structuring element. Lastly, regions with less than 5 pixels were removed. Each region was then manually inspected to determine whether it was a neuron cell body.

To quantify signal colocalization between the eYFP and the PVALB channels, we computed the mean signal intensity of the corresponding soma region in the PVALB channel. The intensity is expressed as a z score, calculated by subtracting each pixel by the mean and then dividing by the standard deviation of all pixels in the image, which allowed us to compare across images. The result is a distribution of intensity values for the soma regions. To determine the threshold at which we can assign soma has having an above background intensity level, we calculated the mean background intensity of its surrounding area (one soma width for each cell, Fig. 3A). We defined the background intensity threshold as the local background PVALB signal intensity corresponding to 5% chance specificity (Fig. 3B pink lines, pink filled pink circle). Any intensity above this threshold is defined as an above background PVALB signal intensity. We then looked up the soma PVALB intensity distribution at this threshold to derive the colocalization specificity (Fig. 3B magenta lines, red filled circle).

### Slice physiology

Rats were anaesthetized with pentobarbital (100 mg/kg) and perfused intracardially with a glycerol-based aCSF (in mM: 252 glycerol, 2.5 KCl, 1.25 NaH_2_PO_4_, 1 MgCl_2_, 2 CaCl_2_, 25 NaHCO_3_, 1 L-ascorbate, and 11 glucose, bubbled with carbogen), and then 300 μm coronal brain slices containing PFC or dSTR were cut in the same solution. Slices recovered at 32 °C in carbogen-bubbled aCSF (containing, in mM: 126 NaCl, 2.5 KCl, 1.2 NaH_2_PO_4_, 1.2 MgCl_2_, 2.4 CaCl_2_, 18 NaHCO_3_, 11 glucose, pH 7.2–7.4, mOsm 302–305) for at least 30 minutes before experiments, with 1 mM ascorbic acid added just before the first slice; experiments were performed in the same solution at 32-34°C. Action potential firing was recorded using Clampex 10.1 or custom routines in IgorPro (Wavemetrics) and an Axon 700A/700B patch amplifier (Molecular Devices). Recordings utilized a K-methanesulfonate based internal solution (in mM: 130 KOH, 105 methanesulfonic acid, 17 HCl, 20 HEPES, 0.2 EGTA, 2.8 NaCl, 2.5 mg/ml Mg-ATP, 0.25 mg/ml GTP, pH 7.2–7.3, ~290 mOsm) or a K-gluconate-based internal (in mM: 9 HEPES, 113 K-gluconate, 4.5 MgCl_2_, 0.1 EGTA, 14 Tris-phosphocreatine, 4 Na_2_ATP, 0.3 Tris-GTP, 10 sucrose, pH 7.2-7.25, ~290 mOsm). Firing was elicited using depolarizing current steps in current-clamp mode, and *Pvalb*+ cell activity was stimulated by ChR2 activation by a 470 nm or 480 nm LED (1 ms pulse width). Cells at ~ −80 mV were depolarized with ~150 pA current step for 500-530 ms.

### *In vivo* optogenetic tagging

*Pvalb^Cre^* rats (n=2) were microinjected with 0.5 μl of AAV5-EF1a-DIO-ChR2(H134)-eYFP virus in dSTR at three locations along a dorsal-ventral trajectory (AP: +0.8, ML: +3.90, DV: −3.0, −4.0, −5.0 from brain surface). In the same surgery, custom drivable optrode bundles, consisting of 16 tetrodes (12.5 μm NiChrome wire, Kanthal) epoxied around a 200 μm core optic fiber, were implanted above dorsolateral striatum (AP: +0.8, ML: +3.8, DV: −2.0 from brain surface). Recording sessions consisted of 90-120 minutes of baseline recording followed immediately by a laser stimulation protocol that delivered light pulses of varying width and frequency. The optrode bundles were driven down into fresh tissue by more than 80 μm at the end of each recording session. Neural data were recorded wideband at 30,000 samples/s with Intan Technologies digital headstages and acquisition board. Single units were clustered offline using automated sorting (MountainSort) and manually inspected in MATLAB.

### Photometry

*Pvalb^Flpo^* rats (n=3) were injected bilaterally with 250 nl of AAV-DJ-Syn-FRT-rev-GCamp6f in PFC and implanted with fiber-optic cannulas (400 μm core, 430 μm outer, NA 0.48, Doric Lenses Inc.). Recordings were done from 4 weeks after implantation. For each rat, the hemisphere with the strongest signal was chosen for analysis. The Doric Lenses two color photometry system was used to provide excitation and record emission. The Tucker-Davis Technologies RZ5D BioAmp processor and Synapse Suite were used to control excitation and data acquisition. GCaMP6f emission from 405 nm (isosbestic point) and 470 nm (calcium-dependent) excitation were recorded. 405 nm excitation was modulated at 330 Hz and 470 nm excitation at 210 Hz. The output excitation intensity was 20-40 μW. Rats were recorded for 3 hours in the home cage and allowed to move freely. Video of the rat, recorded at 30 frames per second, was used for automated reconstruction of the animal’s position and movement speed using Trodes camera module (https://bitbucket.org/mkarlsso/trodes/).

A previously described method [47] was used to process the photometry data. The signals were first filtered with a 5 Hz lowpass filter to remove fluctuations beyond the temporal dynamics of calcium indicator. To subtract the non-calcium-dependent background fluctuations from calcium dependent fluctuations in the signal, we first estimated the background fluctuations based on the 405 nm signal. We smoothed the 405 nm signal (405 nm_smoothed_) using a 5 s moving-average window. This background was then scaled to the 470 nm signal using a least squares linear fit to derive the scaled 405 nm signal (405nm_470nm scaled_). The ΔF/F for the 470 nm signal is (470 nm – 405 nm_470nm scaled_)/405 nm_470nm scaled_. The ΔF/F for the 405 nm signal is (405 nm – 405 nm_smoothed_)/405 nm_smoothed_.

Variation in photometry signal amplitude as a function of speed was quantified as the coefficient of variation (CV), which is the standard deviation of the signal in a 5 s interval expressed as percentage of the mean in the same 5 s interval. Since the background subtracted signals have a mean of 0, we first needed to rescale the data back to the original range by adding back the mean of the background signals (470 nm – 405 nm_470 nm scaled_) + mean(405 nm_470 nm scaled_) or (405 nm – 405 nm_smoothed_) + mean(405 nm_smoothed_).

To compare the strength of correlation between CV and speed, we used linear regression to estimate the slope of the line of best fit between the CV and speed for each animal and recording session. We then compared the slopes for the 470 nm and 405 nm signals using a Wilcoxon rank sum test.

